# Transcriptome-guided annotation and functional classification of long non-coding RNAs in *Arabidopsis thaliana*

**DOI:** 10.1101/2022.04.18.488676

**Authors:** Jose Antonio Corona-Gomez, Evelia Lorena Coss-Navarrete, Irving Jair Garcia-Lopez, Jaime Alejandro Pérez-Patiño, Selene L. Fernandez-Valverde

**Affiliations:** Unidad de Genómica Avanzada, Langebio, Cinvestav, 36824 Irapuato, Guanajuato, México

**Keywords:** Long non-coding RNAs, *Arabidopsis thaliana*, WGCNA

## Abstract

Long non-coding RNAs (lncRNAs) are a prominent class of eukaryotic regulatory genes. Despite the numerous available transcriptomic datasets, the annotation of plant lncRNAs remains based on dated annotations that have been historically carried over. We present a substantially improved annotation of *Arabidopsis thaliana* lncRNAs, generated by integrating 224 transcriptomes in multiple tissues, conditions, and developmental stages. We annotate 6764 lncRNA genes, including 3772 that are novel. We characterize their tissue expression patterns and find 1425 lncRNAs are co-expressed with coding genes, with enriched functional categories such as chloroplast organization, photosynthesis, RNA regulation, transcription, and root development. This improved transcription-guided annotation constitutes a valuable resource for studying lncRNAs and the biological processes they may regulate.

## Background

Long noncoding RNAs (lncRNAs) are transcripts greater than 200 nt with little or no coding potential [1–4]. In contrast to the coding genes, they are smaller, have fewer exons, and have lower expression levels than their protein-coding counterparts [1,3,5–9]. In addition, they often have tissue- and cell-specific expression patterns [1,4,7–9]. lncRNAs have been widely studied in vertebrates. However, few plant lncRNAs have been experimentally characterized to date [10–31].

The available studies on lncRNAs in plants reinforce functional similarities originally observed in animals, including modulation of chromatin topology, miRNA levels (miRNA sponges), precursors of small RNA, and acting as a scaffold for the formation of protein complexes [11,13,32–34]. Plant lncRNAs also participate in the response to biotic and abiotic stresses and environmental stimuli such as bacterial infection [19], salinity [20], drought [25], cold [10,31], nutrient stresses [13,35,36], light [11,18], and heat [26]. They also play a role in reproductive development [10,12,31], growth and development [14,21], chromosome modification [11,22] and the regulation of small RNA abundance via target mimicry [13,18,37]. All the functions mentioned above have in common the interaction of a lncRNA with some other biomolecule (RNA, DNA, or protein).

The search of lncRNAs in plants has resulted in numerous reference annotations. For example, in *A. thaliana*, lncRNAs have been identified and annotated multiple times in competing databases [3,38–43]. Two of the most popular long intergenic non-coding RNAs (lincRNAs) and natural antisense lncRNAs (NATs) reference annotations were generated using 200 *A. thaliana* tiling array datasets and four baseline transcriptomes to annotate all identifiable lincRNAs [4] and a reference annotation for NATs was generated using sense and antisense strand-specific RNA sequencing from 12 strand-specific root transcriptomes [4,44] sequenced in the now discontinued SOLiD sequencing platform [45]. Both of these annotations are now outdated because first, tiling arrays only provide partial information on lncRNA position and expression and can only be used to annotate lincRNAs; second, the SOLiD platform had several problems with decoding when errors occurred during sequencing, as well as with palindromic regions [45]. Moreover, these studies used only four transcriptomes (in the case of lincRNAs), or transcriptomes exclusive to a single tissue (root in the case of the NATs) which limited their capacity to identify a complete suite of lncRNAs, particularly because most of these molecules are expressed in a tissue-specific fashion [9,46].

Several databases store and classify plant lncRNAs [3,38,39,41]. Among these, we wish to highlight the CANTATAdb v2.0 database, which contains 4080 lncRNA genes [41]. The annotations in CANTATAdb are based on ten *A. thaliana* transcriptomes and a robust annotation methodology, including identifying lncRNAs using the Coding Potential Calculator (CPC) [47]. Another important database is GreeNC [38,48], which also uses a predictive annotation through CPC to identify lncRNAs in different species based on transcripts available in Phytozome [49] and ENSEMBL [50], including 2752 genes in *A. thaliana*. In addition, it classifies lncRNAs that can function as miRNA precursors [38]. The most widely used lncRNA reference annotation is Araport11 [40]. Araport11 has 3559 lncRNA genes (2444 lincRNAs and 1115 NATs) [40]. While coding gene annotations in Araport11 arise from the integrative annotation pipeline analysis of 113 RNA-seq experiments on different tissues from plants grown under various conditions, the lncRNAs annotated in Araport11 arise from various sources. In particular, it combines the annotations mentioned above of lincRNAs from [4,44] and the NAT annotations from [44] with lncRNAs well annotated in literature (e.g., *FLINC* and *COOLAIR*) [12,31]. Thus, the lncRNA annotation process in Araport11 was nowhere nearly as strict as their approach to annotating protein-coding genes.

Despite these multiple available sources of annotated plant lncRNAs, few of them have been experimentally characterized or assigned a possible function. A commonly used approach to assign a biological function to lncRNAs is the so-called “guilt-by-association” strategy [51,52]. This involves generating gene co-expression networks and their subsequent functional annotation to assign potential biological functions to lncRNA genes [51,52]. Co-expression networks represent the similarity between the expression patterns of different genes in a set of conditions, developmental stages, and tissues [53]. Genes co-regulated in a wide array of biological conditions are likely controlled by the same regulators or may participate in the same or related biological function or process [52,54–56]. This idea underlies “guilt-by-association” approaches, as lncRNAs can be assumed to work concurrently with the genes it is expressed with, and it is thus preemptively assigned the functions of the genes within its co-expression group. For this approach to work, multiple transcriptomes of the same organism in different stages of development, tissues, and various types of stress are required [53,57,58]. The more transcriptomes used, the better the statistical significance of the co-expression relationship between genes becomes. Furthermore, the diversity of transcriptomes makes it possible to identify specific networks for a condition or tissue and general networks [54]. In plants, co-expression networks have been successfully used for the identification of functions in both coding genes [59–63] and, more recently, in lncRNAs [6,64–69].

To address the need for a better annotation of lncRNAs in *A. thaliana, we* leverage the numerous publicly available RNA-Seq datasets to carry out a comprehensive reannotation of lncRNAs in *A. thaliana*. We reanalyzed 220 publicly available RNA-Seq datasets, in addition to four seedling transcriptomes generated in-house. Furthermore, we integrate these better annotated and expanded lncRNAs within gene coexpression networks, which enable us to identify potential functions.

## Construction and content

### Publically available transcriptomes used

We selected 220 publicly available transcriptomes using the following criteria: 1) a minimum of 0.5 gigabases (GB) per transcriptome, and 2) generated in a condition, tissue, or developmental stage of wild-type *Col-0 A. thaliana*. These included: embryo, seed, hypocotyl, cotyledon, root tip, apical shoot meristem (ASM), seedling, root, plant callus, petiole, leaf, carpel, flower pedicel, petal, pollen, sepal, stamen, flower, stem internode, stem node, septum, valve, whole adult plant and conditions such as cold, heat, salinity, drought, blue light, red light, limited phosphate, limited iron and presence of abscisic acid (ABA). All transcriptomes were downloaded as raw reads from Gene Expression Atlas (GEA) [70] and Gene Expression Omnibus (GEO) [71]. Each dataset is described in detail in Table 1. Additionally, we generated four transcriptomes from the aerial part and roots of *A. thaliana* 8 day post-germination seedlings (see details below), totaling 224 transcriptomes (Dataset S3).

**Table 1.**
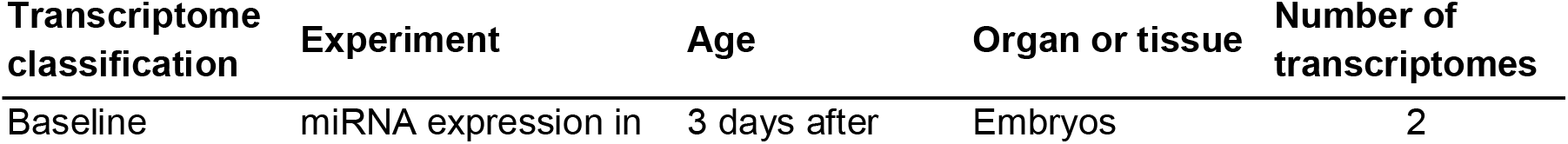
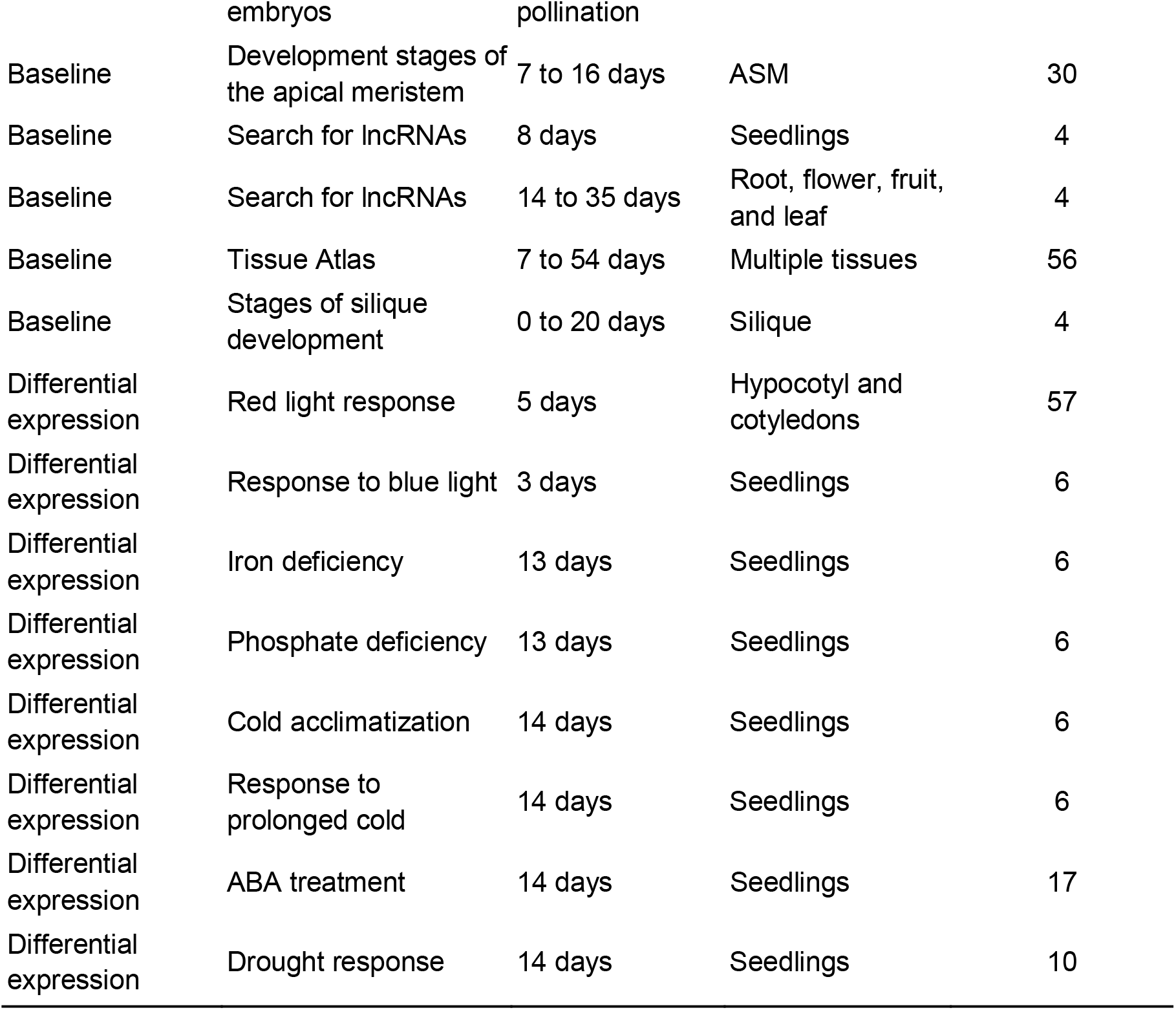
Summary of transcriptomes used for lncRNA annotation

### In-house transcriptome generation

Seedlings were grown *A. thaliana* in Murashige and Skoog (MS) solid medium within growth chambers under conditions of long days (21 °C, 16 / 8h photoperiod cycles), approximately for 8 days. The aerial part (shoot) and roots were collected separately, with two biological replicates for each organ (fully open cotyledons and 2 rosette leaves greater than 1 mm long). Total RNA was extracted using TRIzol (Invitrogen, 15596018), and according to company specifications, samples were DNase I treated using TURBO™ DNase (Invitrogen, AM2238). The quality and concentration of the samples were measured using the NanoDrop 2000C spectrophotometer (Thermo Fisher Scientific Inc). The integrity of the RNA was verified using a 1.5% agarose gel, and the cDNA was synthesized using the NEBNext Poly (A) mRNA Magnetic Isolation protocol (NEBNext, E7490S). The libraries were prepared using the NEBNext Ultra II Directional RNA library kits (NEBNext, E7760S) and NEBNext Multiplex oligos for Illumina (SET 1) (NEBNext, E7335). The libraries were sequenced using the Hi-Seq X from Illumina, using 2×150 nt (PE150). The depth and characteristics of these libraries are summarized in Table S1.

### Filtering, assembly, and quantification of transcripts across all transcriptomes

We assessed the quality of all transcriptomes using FastQC v0.11.2 [72] and MultiQC v1.0 [73]. Low-quality reads and adapters were removed using Trimmomatic v0.32 (HEADCROP:10-5 LEADING:5 SLIDINGWINDOW:4:15 MINLEN:30-60) [74]. All quality filter reads were aligned to the *A. thaliana* TAIR10 [75], using STAR v2.7.2.b (--alignMatesGapMax 120000) [76]. The resulting alignments were assembled using StringTie v1.3.4 (-f 0.3 -m 50 -a 10 -j 15 -c 2.5) [77], using the Araport11 annotation as a reference [40]. The resulting transcripts were joined using the merge function (-c 2.5 -f 0.3) of the StringTie v1.3.4 program [77]. Transcript counts were obtained using Kallisto v0.44.0 (parameters for single-end transcriptomes: --single -t 8 -l (40, 67, 80) -s (5, 10, 20); parameters for paired-end transcriptomes: default) [78].

### lncRNA identification

To identify the lncRNAs, we first generated the amino acid sequence for all transcripts using TransDecoder v5.3.0 [79]. We then applied nine sequential filters based on previous studies [5,9] (see Fig. S1). We refer to this process as the Strict Method (SM). First, (1) we selected all autosomal transcripts ≥ 200 nt using the infoseq program of EMBOSS v6.6.0 [80]. We eliminated sequences whose translated ORF or nucleotide sequence had homology to proteins in the Uniprot database [81] as measured by the (2) blastp (e-value ≤ 1e-6) or (3) blastx (e-value ≤ 1e-6, strand=“plus”) program, respectively [82]. We subsequently removed sequences with (4) identifiable protein domains found in the base of Pfam (v33.0) [83] using the HMMER v3.1b2 program [84] (e-value ≤ 1e-6) or (5) with identifiable signal peptides using signalP v4.1 [85] (D-cutoff: 0.45). For any reminder sequences, (6) we removed those that had an ORF > 100 aa using the program getorf of EMBOSS v6.6.0 [80]. We did an additional filtering step of all sequences with homology to non-redundant proteins (nr) annotated in the NCBI database [85,86] using BLASTx [82] (evalue ≤ 1e-6, strand = “plus”). For each remaining transcript, we identified the best blast hit against the ‘nr’ database with a percentage of identity above 70% (pident ≥ 70.000). For each best hit, we used the blastdbcmd function [82] to obtain the information related to the ID. The transcripts annotated in NCBI as: “hypothetical protein” (in Refseq), “similar to” (NCBI’s annotation pipeline), “putative protein”, “unknown (unknown protein, unknown, partial, unknown)”, “predicted protein” and “unnamed protein product” [87] were retained. tRNAs and rRNAs were identified using infernal v1.1.2 [88] and the covariance models in the Rfam database [89]. We additionally compared sequences with tRNAs and rRNAs reported in *A. thaliana* using BLASTn [82] (evalue ≤ 1e-6, strand = “plus”). All sequences identified as tRNAs or rRNAs were discarded. Finally, we eliminated transcripts with introns > 6000 bp.

After filtering, we manually reviewed transcripts classified in Araport11 [40] as coding proteins or genes and in our annotation as lncRNAs. This manual review consisted of verifying if these genes had annotation as functional proteins or annotated domains; in these cases, the lncRNA was discarded; if it was a hypothetical or not described protein, the lncRNA was retained. Thus, all sequences that passed this final review constituted the final set of SM lncRNAs.

### Classification of lncRNAs by genomic position

LncRNAs are generally classified by their positional relationship to other genes. We used the following non-overlapping categories, based on the GENCODE annotation [1]:

#### Intergenic lncRNAs (lincRNAs)

lncRNAs found in intergenic regions.

#### Natural antisense lncRNA (NAT)

lncRNAs that totally or partially overlap an exon of another gene in the complementary chain.

#### Sense-exonic lncRNAs

lncRNAs that totally or partially overlap the exon of another gene with the same direction of transcription (transcribed from the same DNA strand).

#### Intronic lncRNAs

lncRNAs found within the intron of another gene without overlapping any of its exons, including those on the same chain or complementary to the superimposed gene.

It is worth mentioning that all the isoforms of the overlapping gene are considered for all these categories. To know with which genes our lncRNAs overlap, we used the annotation of Araport11 [40] and BedTools intersectBed (sense_exonic lncRNAs [-wo -f 0.1 -s], NAT [-wo –f 0.1 -S], intronic [-wo -f 1] and lincRNAs [-wo -v]) [90]. Finally, all final annotations were inspected by visualizing them in the UCSC Genome Browser [91].

### Comparisons with other lncRNA databases

The 6764 genes annotated as lncRNAs by the SM were compared with the 2752 genes in GreeNC (v1.12) [38], 4080 genes in CANTATAdb (v2.0) [41] and 3559 genes in Araport11 [40]. We compared the coordinates between these databases using the intersectBed program (-wo –s -f 1 -F 1) from the BedTools toolkit [90]. We visualized all lncRNA annotations in the UCSC Genome Browser and corroborated the gene assignment for each lncRNA transcript. We summarized these comparisons using the VennDiagram [92] and UpSetR packages [93] in R.

### Quantification of lncRNAs by tissue and stage

The transcriptomes were divided into tissue and developmental stage categories based on their age and tissue of origin. Notably, some categories are not *bona fide* tissues (e.g. whole plant, seedlings). However, these were considered their own category as these transcriptomes can be readily differentiated from others. All the transcriptomes were classified into five developmental stages based on the classification by [94] (Fig. 2b). The first two stages belong to the vegetative phase and include: seed germination (Stage 1, 3 to 5 days old) and leaf development (Stage 2, 6 to 25 days old); the rest of the stages are part of the reproductive phase, ranging from the presence of the first inflorescence (at 26 days old) (Stage 3, 26 to 29 days old), flower production (Stage 4, 30 to 47 days old), to the generation of siliques (Stage 5, 48 to 51 days old) (Table 1-transcriptomes, Fig. 2b).

**Figure 1.**
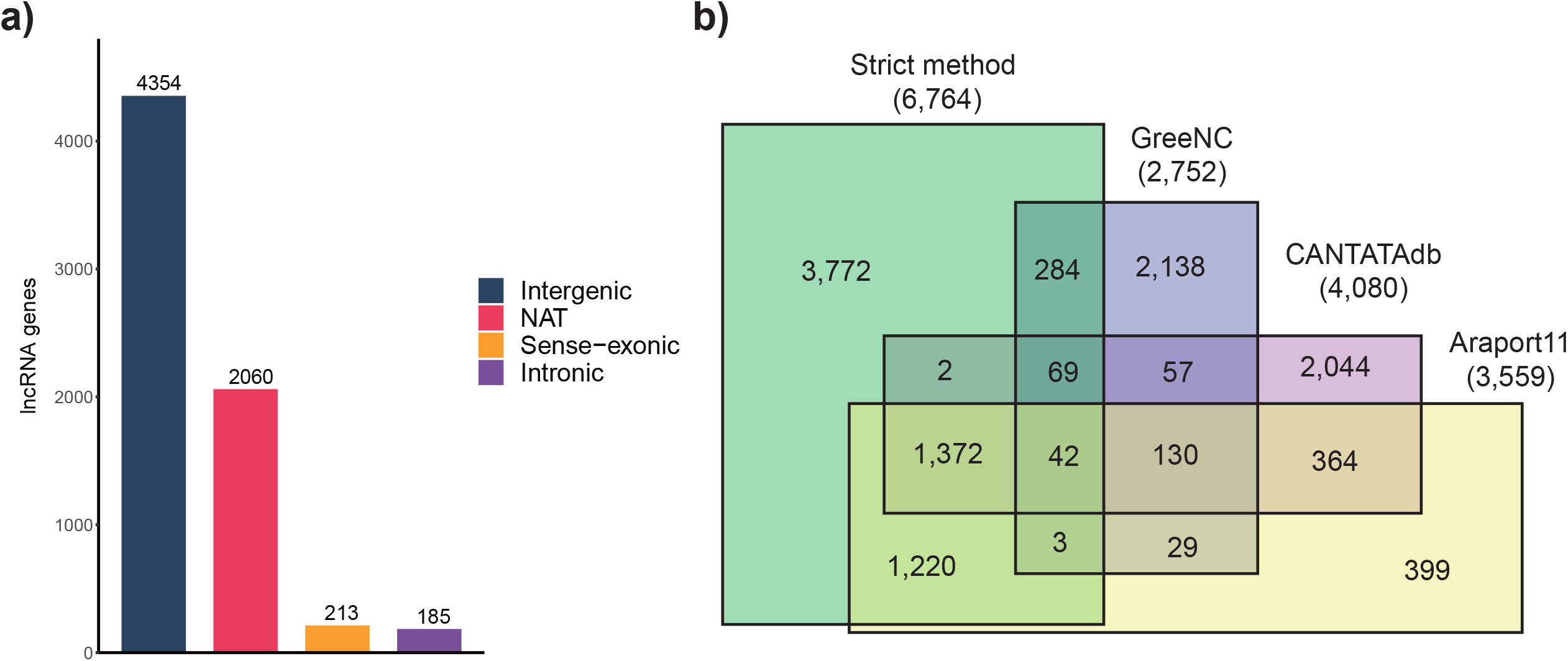
Annotation of lncRNA genes and comparisons with other plant lncRNA databases. a) Distribution of the 6764 lncRNA genes predicted by the SM. b) Venn di agramcomparing the SM (green) with the databases GreeNC (purple), CANTATAdb (pink) and Araport11(yellow) where the lncRNAs have been annotated from *A. thaliana*. SM=Strict Method.

**Figure 2.**
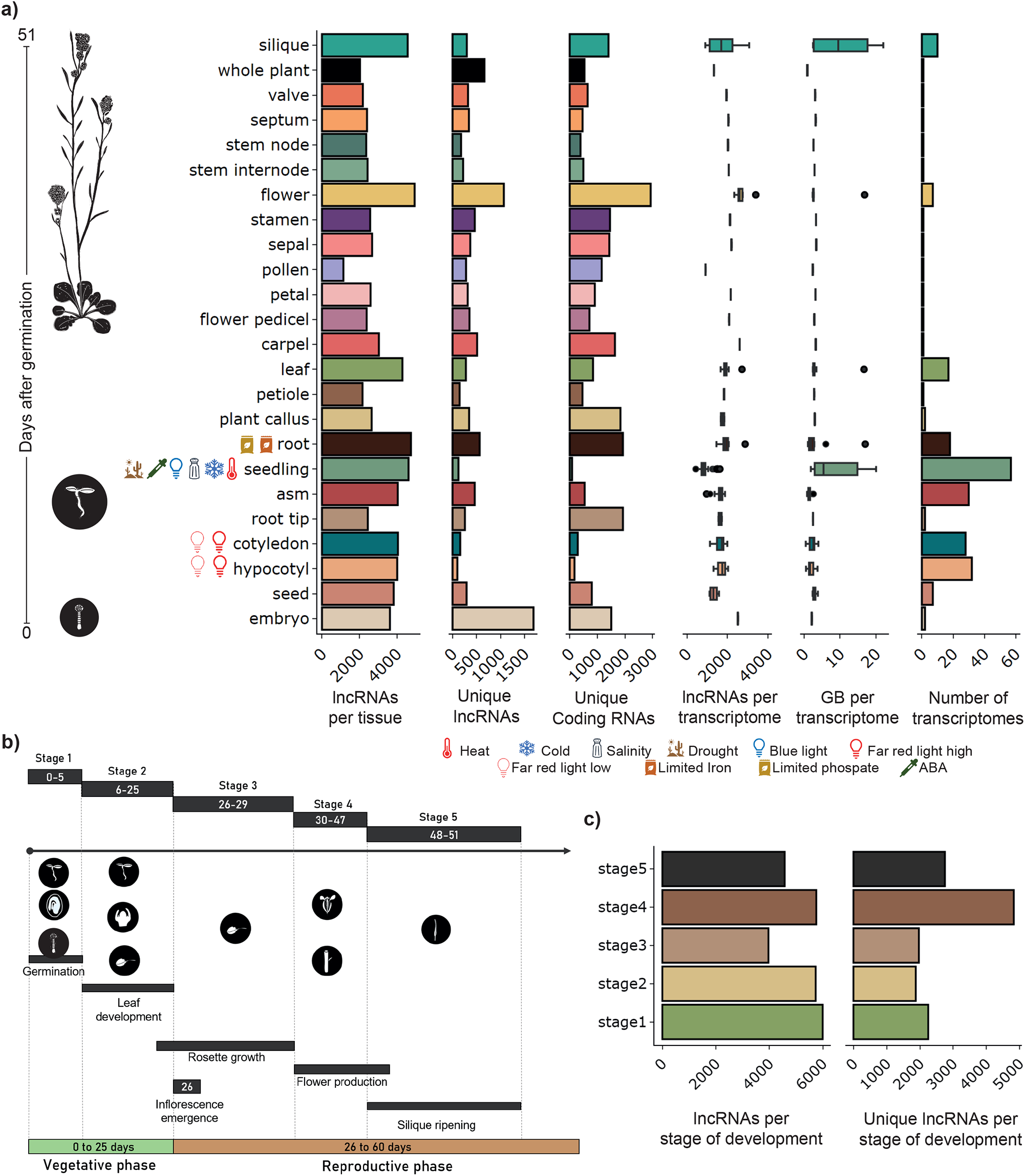
Expression of lncRNAs by tissue and life stage. a) Histograms of the number from left to right of (x-axis): lncRNAs per tissue, unique lncRNAs, unique coding RNAs, followed by box-plots displaying the number of lncRNAs per transcriptome, GB per transcriptome and number of transcriptomes found in each tissue and life stage studied (y-axis), ordered by the mean number of days after germination. b) The tissues were grouped into five life stages of *A. thaliana* using their position within the *A. thaliana*’s developmental progression. c) Number of lncRNAs per life stage. The histogram contains the number of lncRNAs identified in each of these stages, while the second histogram contains the number of unique lncRNAs for each of these stages.

To identify lncRNAs specific to a tissue or stage of development, we calculated the value of SPM (Specificity Measure) based on the work methodology described in [95]. The SPM value of a gene is obtained by dividing its mean transcripts per million (TPM) value in a category with the sum of the TPM values in all categories, resulting in a value from 0 to 1 where genes that are tissue or stage-specific have values close to 1. Only the top 5% values close to 1 in all transcriptomes were considered tissue-specific or developmental stage-specific (Fig. S2).

### Generation of coding and non-coding gene co-expression networks

To determine the possible functions of lncRNAs, we used a *guilty-by-association* approach. This approach identifies enriched functional annotations of protein-coding genes co-expressed with the lncRNAs, which allows inferring the biological processes in which these lncRNAs may be involved. The co-expression network was built using the WGCNA tool [96] based on the table of raw counts for the full transcriptome normalized using the variance stabilizing transformation (VST), part of the DESeq2 package [97]. The adjacency function was weighted by the power of correlation between the different genes, and the law of free-scale networks determined the parameter β. To ensure that the average connectivity of the network was continuous, we chose a value of β = 12, which is the lowest value for which the unscaled topology index curve remains stationary. From this point on, we will refer to the groups of co-expressed genes as co-expression modules or simply modules, following the nomenclature used by the WGCNA program [64]. The network was of type signed with a bicor correlation (biweight midcorrelation)

and the option of separate modules (unmerged) with a minimum module size of 50 genes. The expression profiles were represented by their main component (module eigengene). An eigengene is the first right-singular vector of the standardized gene expression [98] that serves as a summarized representation of the expression of all genes in each module. To identify the functions associated with each co-expressed module, we performed an enrichment analysis of Gene Ontology (GO) categories using topGO [99] for the Biological Process (BP) ontology. Finally, we used a Fisher test (qval.bh < 0.01) to assess the significance of the enrichment of GO categories.

### Genome browser

All lncRNA annotations were uploaded to the UCSC Genome Browser as a track for visualization [91]. The coordinates of all lncRNAs genes are available in the Dataset S1 in bed12 format.

## Results

Using the SM, we identify 6764 lncRNA genes (7070 transcripts). These included 4354 lincRNAs (4549 transcripts), 2060 NATs (2133 transcripts), 213 sense-exonic (248 transcripts), and 185 intronic (187 transcripts) (Fig. 1a, Dataset S1), 78 intronic lncRNAs had no transcriptional orientation (sense) as they were identified in single-end transcriptomes only. Furthermore, 33 lncRNA genes (46 transcripts) were categorized as both NATs and sense-exonic due to the position of the lncRNA flanked by both sense and antisense coding genes in the DNA strand. These were manually verified to ensure they were not extended 3’ UTRs of overlapping protein-coding genes. Additionally, 15 genes had isoforms belonging to different categories (Dataset S2).

As expected, the identified lncRNAs have fewer exons per transcript (median 1; average 1.23) (Fig S3a) than coding genes (median 4; average 6). Furthermore, their mature transcripts are smaller (average 437.3 nt) than that of their coding counterparts (average 1799 nt) (Fig. S3b). These characteristics coincide with what has been previously observed in animals [5,7,8,100], flies [101] and other plants [68,102–105].

The total of lncRNAs annotated by the SM (6764 genes) outnumbers the most prominent databases in *A. thaliana*: GreenNC (v1.12) has 2752 genes (3008 transcripts) [38], CANTATAdb (v2.0) 4080 genes (4373 transcripts) [41] and Araport11, 3559 genes (3970 transcripts) [40]. A comparison with these databases revealed that 3772 lncRNAs genes in our annotation are novel and have not previously been reported in any of these databases (Fig. 1b); the new lncRNAs were categorized into 2326 lincRNAs (2454 transcripts), 1218 NATs (1227 transcripts), 111 sense-exonic (124 transcripts) and 145 intronic (146 transcripts). These new lncRNAs represent a 93.08% (2275 over 2444) increment in the number of lincRNAs in lincRNAs and a 134.70% (1502 over 1115) increment in NATs, with respect to the Araport11 database. Additionally, we find that 398 lncRNA genes of lncRNAs are shared between our annotation and the GreeNC database [38], 1485 with CANTATAdb [41], and 2637 with Araport11 [40], being the Araport11 database the one with the best agreement with our data; our annotation contains approximately 74.09% (2637 over 3559) of the lncRNAs annotated in Araport11 (Fig. 1b).

Surprisingly, only 130 lncRNAs are shared between GreeNC, CANTATAdb, and Araport11 databases, and there are only 42 lncRNAs shared among the four annotations (Fig. 1b). It is important to note that there are likely other lncRNAs in *A. thaliana* that are not identified in our analysis, since not all conditions, tissues, and developmental stages have been surveyed using RNA-Seq. However, our annotation is the first to take advantage of most of the transcriptomic data available for this species, ensuring that the sequences obtained are only those of expressed lncRNAs. This, combined with a robust annotation method, avoids redundancy with other types of transcripts that are not lncRNAs.

Interestingly, when comparing our annotation to Araport11, we observe that our annotations were not always in the same biotype classification. The most concordant classification between both annotations was among lincRNAs, where 1747 lincRNA genes correspond to the same annotation (Fig. S4). However, several lncRNAs identified in our annotation are not classified as lncRNAs in Araport11: 288 lncRNA genes (265 lincRNAs, 12 NATs, 6 sense-exonic, 4 lincRNA-NAT, and 1 sense-exonic lincRNA) are annotated in Araport11 as “novel transcribed region”, and 388 as “coding genes’’ (217 lincRNAs, 110 NATs, 43 sense-exonic, 13 NAT-sense-exonic, 3 sense exonic lincRNAs, 1 lincRNA-NAT and 1 intronic) (Fig. S4).

We were particularly interested in these 388 lncRNAs classified as “coding genes” in Araport11. We manually reviewed these annotations and concluded these are, in fact, lncRNAs that are erroneously annotated as “coding genes’’ in Araport11. Among these, we found *IPS1* (Induced by Phosphate Starvation 1, AT3G09922), a lncRNA with a mimicry target function for microRNA miR399 in the absence of phosphate [13]. Another erroneously classified lncRNA was *IPS1’*s paralog *At4* (AT5G03545) [37], which is functionally redundant to *IPS1*. Both of these lncRNAs have been previously experimentally validated and found to be conserved across several plant species [106–109]. Similarly, the *lncRNA APOLO* (AUXIN-REGULATED PROMOTER LOOP, AT2G34655) [16] is annotated as a protein-coding gene. We also found multiple lncRNAs erroneously annotated as snoRNAs, novel transcribed regions, and other RNAs, including the experimentally validated lncRNAs: *HID1* (HIDDEN TREASURE 1, AT2G35747) [11], *MARS* (MARneral Silencing, AT5G00580) [22], and *DRIR* (Drought-induced RNA, AT1G21529) [20], respectively (Table 2).

**Table 2.** lncRNAs erroneously annotated in Araport11

| Annotated lncRNAs | Araport11 annotation | Araport11 gene IDs | Other lncRNAs databases | Biological role | Biological condition | Experimental validation |
| --- | --- | --- | --- | --- | --- | --- |
| *IPS1* | protein-coding | AT3G09922 | GreeNC, EVLncRNAs and PNRD | Phosphate homeostasis | Phosphate deficiency | [13] |
| *At4* | protein-coding | AT5G03545 | GreeNC, EVLncRNAs and PNRD | Phosphate homeostasis | Phosphate deficiency | [37] |
| *APOLO* | protein-coding | AT2G34655 | GreeNC, EVLncRNAs and PNRD | Auxin-controlled development | Auxin | [16] |
| *HID1* | snoRNA | AT2G35747 | EVLncRNAs | Photomorphogenesis | Continuous red light | [11] |
| *DRIR* | other RNA | AT1G21529 | NONCODE (NONATHT00017 2.1), PNRD (NONATHT00017 2) and EVLncRNAs | Stomatal closure | Drought, salinity and ABA | [20] |
| *ELENA1* | other RNA | AT4G16355 | GreeNC and CANTATAdb, NONCODE (NONATHT00289 9.1.1) | Defense | *Pseudomonas syringae* pv.tomato DC3000 | [19] |
| *TAS1a* | other RNA | AT2G27400 | GreeNC | Acclimation and freezing tolerance | Cold | [141] |
| *MARS* | novel transcribed region | AT5G00580 | - | Germination | Response to ABA | [22] |
| *BLIL1* | NAT | AT1G26218 | CANTATAdb, GreeNC and NONCODE (NONATHT00009 2.1) | Photomorphogenesis | Blue light | [18] |
| *COOLAIR* | NAT | AT5G01675 | EVLncRNAs and CANTATAdb | Flowering time | Cold | [31] |
| *FLORE* | NAT | AT1G69572 | CANTATAdb, GreeNC and NONCODE (NONATHT00083 4.1.1) | Circadian rhythms | Long days, short day and 12h light/12h dark | [24] |
| *FLINC* | lincRNA | AT1G08103 | PlncDB (ID=ncrna9858) | Flowering time | At 16 °C | [12] |

In addition to these categories, we identified numerous lncRNAs that were annotated as transposable elements (92, reclassified as 91 lincRNAs and 1 NAT), other RNA (83: 77 lincRNAs, 3 lincRNA-NAT, 1 NAT, 1 sense-exonic and 1 sense-exonic lincRNA), pseudogenes (48: 43 lincRNAs, 2 NAT, 2 sense-exonic and 1 NAT lincRNA), snoRNA (5: 5 lincRNAs) and snRNA (1: 1 NAT) (Fig. S4). Finally, we found 3222 lncRNA genes that are not shared between Araport11 and our annotation. These lncRNAs comprise 1885 lincRNAs, 1117 NATs, 143 intronic, 67 sense-exonic, and 10 genes shared between NATs and sense-exonic (9 genes) and intronic and sense-exonic (1 gene) (Fig. S4). These last 10 genes had two annotations due to having isoforms belonging to two different categories. This comparison shows that the annotation of lncRNAs in Araport11, one of the most prominent reference databases for *A. thaliana*, has significant inaccuracies that are resolved in our annotations, resulting in an improvement in the classification of lncRNA genes.

It is worth noting that within our annotation, 48 lncRNA genes (120 transcripts) have an ambiguous annotation, as they are simultaneously annotated as NAT and sense-exonic (33 lncRNA genes; 93 transcripts), lincRNAs and sense-exonic (5 genes; 12 transcripts), lincRNAs and NAT (9 genes; 13 transcripts), and intronic and sense-exonic (1 gene; 2 transcripts) (Dataset S2). Specifically, in the case of lncRNAs annotated as NAT sense-exonic, they overlap two different protein-coding genes, thereby acquiring a separate annotation for each gene. Similarly, other lncRNA genes had isoforms in different categories, depending on the genomic location of each isoform.

### Expression patterns of lncRNAs

In addition to annotating lncRNAs, we leveraged the transcriptomic information to explore how lncRNA expression was distributed amongst *A. thaliana* tissues, developmental stages, and conditions (Fig. 2). We found more lncRNAs expressed in flower, root, seedling, and silique (Fig. 2a). LncRNAs were often more abundant in organs with higher cell-type diversity, such as flowers, silique, roots, and leaves (Fig. 2a). This tendency has been previously observed in animals, where organs with more diversity of cell types, such as the brain, express more lncRNAs [110–112]. Reproductive tissues are also known to host a greater diversity of lncRNAs. Similarly, in our data, flowers have more lncRNAs than other organs (Fig. 2a). Interestingly, the number of lncRNAs expressed in the flower is much higher than in its individual parts (stamen, sepal, petal, carpel, and pedicel), further suggesting the high tissue and cell-type diversity of this organ may be due to the multiple tissues that make up this organ. An enrichment of lncRNAs in reproductive tissues has been previously reported in multiple plant species such as soy, corn, and rice [34,113,114] and animal testis [9,57,115,116]. Another category that stands out for its abundant lncRNAs is seedlings (Fig. 2a), composed of a mixture of tissues in a particular developmental stage. As most of the transcriptomes from abiotic stress conditions used in this study were from seedlings, many lncRNAs expressed in response to these stresses are expressed in and thus assigned to seedlings (Fig. 2a). Also, the number of transcriptomes and the sequencing depth in each category correlates positively with the number of lncRNAs found (Fig. 2a).

In terms of development, the vegetative phase (stages 1 and 2) has the highest number of lncRNAs (Fig. 2c), followed by stage 4 (flower development). Developmental stages where tissue differentiation or organ formation occur tend to express multiple lncRNAs in both plants [6,117–119] and animals [8,46,57,111]. Unfortunately, the early stages of tissue differentiation are not represented in our data set, which could help us identify lncRNAs that participate in tissue formation.

### Tissue and stage-specific lncRNA expression

Genes specifically expressed in a particular tissue or stage of development may be important for establishing the identity of that tissue or stage. We found that lncRNAs in *A. thaliana*, as in most organisms, are expressed in a more tissue-specific manner compared to coding genes (Fig. 2a). The embryo and the flower had the highest amount of unique lncRNAs, while the flower and the root expressed more unique coding genes (Fig. 2a). Interestingly, despite not being the most abundant in lncRNAs (Fig. 2a), the embryo has the highest number of unique lncRNAs. Also, the flower expressed most of the unique coding and lncRNAs genes (Fig. 2a). Interestingly, there were many more unique coding genes in the root and almost no unique lncRNAs expressed (Fig. 2a). We did not observe an increase in unique genes in tissues with various stress conditions. Also, most unique lncRNAs are expressed in the reproductive stages of the plant rather than in the juvenile stages (Fig. 2c). Dividing the lncRNAs by biotype, we find that 94.0% (4093 of 4354) of lincRNAs, 88.11% (1815 of 2060) of NATs, 93.4% (199 of 213) sense exons, and 50.27% (93 of 185) of intronic lncRNAs belong to a single tissue or stage, corroborating that lincRNAs display more tissue specificity than NATs. These results indicate a high specificity of lncRNAs in the different tissues and stages.

### LncRNAs with known tissue-specific functions

Some lncRNAs with known functions display a tissue-specific expression pattern related to their function (Table S2). Among these, we find lncRNAs *IPS1* and *At4*, which have functions related to phosphate starvation [13,37], and the lncRNA *MARS*, which is involved in changes of the chromatin conformation in response to ABA [22]; all three of which we find expressed in root tissues. In addition, the lncRNA *FLINC* is expressed in flower and meristem tissues, which is related to the regulation of flowering [12]. On the other hand, the tissue-specific expression of some known lncRNAs does not coincide with what has been previously reported in the literature. For example, this is the case of *HID1*, which is found in the apical meristem, while it has been reported in cotyledons and hypocotyls [11]. Another example of this is *APOLO*, which participates in lateral root development in response to auxin [16,120], and we find more expressed in the petiole.

### Co-expression of lncRNAs with coding genes

To infer a possible function for all annotated lncRNAs, we used a so-called *guilty-by-association* approach. To this aim, we constructed a co-expression network including all coding and non-coding genes using WGCNA [96]. A total of 224 transcriptomes with 34,937 genes were analyzed to construct this network.

We obtained a total of 45 co-expression modules (Fig. 3). 1425 (21%) lncRNA genes were found in 44 of the 45 co-expression modules. Overall, 516 lincRNAs, 746 NAT, 104 sense-exonic, and 59 intronic were co-expressed. Module 1 harbored the most lncRNAs, with 383 of them, primarily NATs (290), followed by lincRNAs (79), 13 sense, and one intronic lncRNA (Fig. 3). According to the functional enrichment for biological processes, this module stood out for processes related to photosynthesis, the organization of chloroplasts, and response to light. The next modules with the highest number of lncRNAs are modules 4 and 3; these modules are related to the processing and transcription of RNA. In total, 91% (40/44) of modules that housed lncRNAs presented functional enrichment for biological processes. Interestingly, 746 novel lncRNAs were co-expressed with coding genes and distributed amongst 40 modules. Modules 1, 3, 4, and 6 had the most newly annotated lncRNAs, most of them NAT lncRNAs (Fig. S5). It is worth noting that most novel lncRNAs (3026, 80.2%) in our annotation were not co-expressed with coding genes.

**Figure 3.**
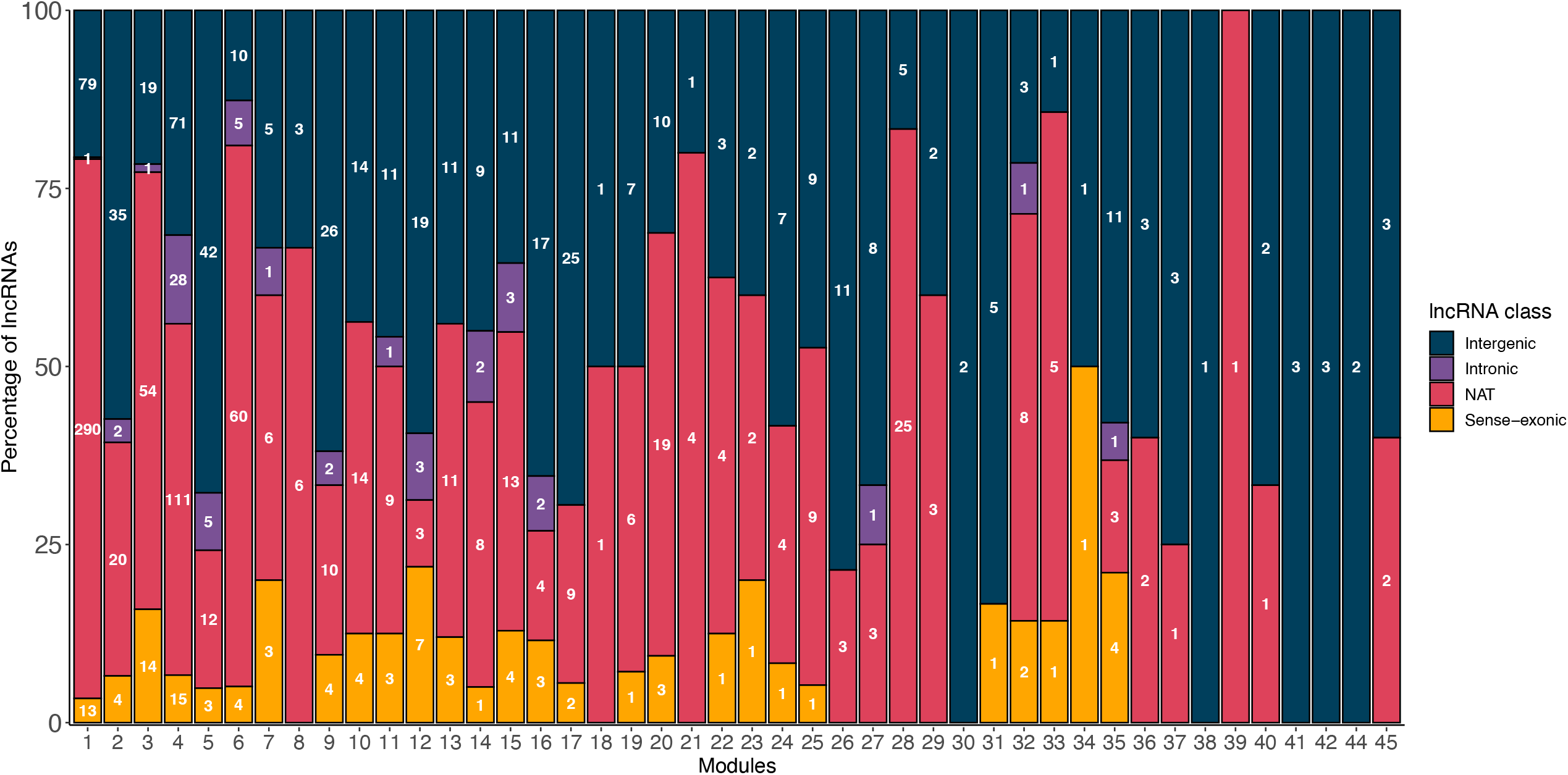
LncRNAs per module. Number of lncRNAs in each co-expression module and classified by biotype (Intergenic (lincRNA) -blue, Intronic -purple, NAT -pink, Sense-exonic -yellow).

We found that the 40 coexpression modules that housed lncRNAs and had functional enrichment could be grouped into 9 categories (Fig. S6 to S14) mainly by their function followed by their eigengene values (representative gene expression) in the different tissues or developmental stages [96]. These functional categories are chloroplast organization and photosynthesis (4 modules with 409 lncRNAs) (Fig. S6), RNA regulation and transcription (4 modules with 375 lncRNAs) (Fig. S7), root development and response to root-related stress (5 modules with 125 lncRNAs) (Fig. S8), protein labeling and transport with (5 modules with 117 lncRNAs) (Fig. S9), cell division (5 modules with 112 lncRNAs) (Fig. S10), lipids and membranes (3 modules with 97 lncRNAs) (Fig. S11), response to pathogens (4 modules with 72 lncRNAs)(Fig. S12), DNA repair (2 modules with 61 lncRNAs)(Fig. S13) and response to stress (5 modules with 17 lncRNAs) (Fig. S14).

We found that the largest number of coexpressed lncRNAs are in the functional category enriched for chloroplast organization and photosynthesis, with positive expression eigengene values in organs related to photosynthesis such as leaves, cotyledons, and hypocotyls. These lncRNAs are divided into four modules (Fig. S6) related to more specific functions such as response to radiation (response to red light, high-intensity light) (module 1), chloroplast and plastid organization (modules 1, 32, and 33), response to cold (modules 1 and 36) and seed, embryo development (modules 32, 33 and 36). The following functional category where we find numerous lncRNAs is related to RNA regulation and its transcription. This category comprises four modules (Fig. S7) with functions such as mRNA metabolic process (module 4), RNA processing (module 3), and regulation of gene expression (modules 16, 17). Genes in these functional categories are most highly expressed in embryos, apical meristem, and plant callus. The expression profile in this functional category is very similar to the function category of cell division (5 modules) (Fig. S10), which has positive expression values in embryos, seeds, apical shoot meristem, and roots. The group of modules with fewer lncRNAs is enriched in genes that participate in the response to abiotic conditions (Fig. S14). However, many modules (modules 1, 2, 5, 6, 14, 19, 23, 36, and 45) are enriched in genes that participate in other stress responses (such as drought and cold). Still, they were classified in other functional groups, such as root development (Fig. S8).

Several functionally characterized lncRNAs belong to specific functional categories. For example, the *DRIR, AT4*, and *APOLO* lncRNAs are found in the group of modules related to root function and stress response. It is known that *DRIR* regulates the closure of stomata in drought [20], *AT4* is associated with the response to phosphate deficiency [121], and *APOLO* is a regulatory lncRNA that directly controls its neighboring gene PID and a many of independent genes by DNA association in response to auxin [16,120]. The functions of these lncRNAs fit what we observed in the functional enrichment of the modules where they are found. In addition to these examples, we have some others in the group of chloroplasts and photosynthesis, such as *FLORE* [24]. This lncRNA has been identified as an important factor in the photoperiod. The lncRNA *FLINC*, identified as a mediator of flowering in response to temperature [12], is found in the group of RNA regulation and transcription functions.

## Discussion

Here, we generated a new and improved annotation of lncRNAs in *A. thaliana*, supported by 224 transcriptome datasets (Dataset S3) obtained from 24 organs (parts of the plant), 11 conditions, and 5 developmental stages (20 timepoints) (Fig. 2). We found 6764 lncRNAs genes (7070 transcripts), including 3772 novel lncRNAs (Fig. 1b). Among our annotated lncRNAs, we identified 58 genes (86 transcripts) of lncRNAs experimentally validated in *A. thaliana* from the EvlncRNAs database [122], which supports our ability to identify functionally relevant lncRNAs by leveraging existing publicly available transcriptomes.

Given our much cohort of transcriptomic evidence, we find few lncRNAs shared with databases such as GreeNC (398 lncRNAs genes shared) [38] and CANTATAdb (1485 lncRNAs genes shared) [41], and about 74.09% of the lncRNAs in Araport11 are found in our annotation [40] Importantly, our curation approach helped us identify several lncRNAs that were erroneously annotated as coding genes including the lncRNA *IPS1*, an experimentally validated lncRNA with multiple target sites for miR399, induced in the absence of phosphate [13]. Another example is its paralog *At4*, which presents functional redundancy with *IPS1* [37]. Although these two lncRNAs are functionally conserved in tomato (*Lycopersicon esculentum L*.) (lncRNA *TPSI1*) [106], *Medicago truncatula* (lncRNA *Mt4*) [107], rice (lncRNA *OsIPS1*) [108] and barley (*HvIPS1*) [109]. Similarly, the well-characterized lncRNA *APOLO*, which regulates lateral root development [16], is annotated as a protein-coding gene. The experimentally characterized lncRNAs *HID1* [11], *MARS* [22], and *DRIR* [20] were also erroneously classified (Table 2). Given the wide usage of the Araport11 database, we recommend a revision of their annotations based on our results.

The hundreds of transcriptomics datasets we used allowed us to analyze the abundance of lncRNAs in the different tissues and the development stages. Our analysis revealed that organs with more cell-type diversity display the highest abundance of lncRNAs in *A. thaliana* (Fig. 2a). This pattern is particularly prominent in organs related to reproduction (flower & silique), similarly to previous reports in multiple animals [9,57,115,116] and plant species [34,113,114].

We find that the depth and the number of the transcriptomes are the experimental factors that most affect our capacity to identify novel lncRNAs in any given sample, similarly to previous annotation efforts in various species [114,123,124]. Thus, we recommend having higher sequencing depth to expedite the discovery of lncRNAs. One limitation of our study is the lack of data from stages where tissue differentiation occurs in the plant, including the flower formation and embryonic stages and the formation of the gametes—surveying these biological conditions would be essential to help complete the catalog of *A. thaliana* lncRNAs and further our understanding of the role of lncRNAs in the formation of plant structures.

In animals, organ formation and differentiation primarily occur in the embryonic stage, while in plants, it occurs not only in the embryonic phase but also in germination and flower development (Fig. 2b). It has been shown that widely expressed and conserved lncRNAs are expressed during tissue development, which has the highest probability of being functional. As the tissue matures, an increasing number of species and organ-specific lncRNAs are more likely to be non-functional [46].

We find that the expression of lncRNAs is more specific than the expression of coding genes. Nearly 90% (6231) of lncRNAs have expression profiles restricted to a specific tissue or stage, while only 55% (16,490) of proteins are specific to a particular tissue or stage. This finding agrees with previous reports in *A. thaliana* [4] and other species [1,9,125,126]. Moreover, most lincRNAs displayed high tissue specificity, while intronic lncRNAs and NATs had low tissue specificity. This is a particularly puzzling observation in the case of NATs, as they generally regulate the gene with which they are associated [127,128]. However, they have higher tissue specificity than their coding counterparts, which generally have less tissue-restricted expression patterns.

We found 1241 co-expressed lncRNAs, which we could associate with our broad functional categories (Fig. 3). However, the functions that we can assign to lncRNAs are limited by our set of transcriptomes; we can only identify enriched biological functions in the tissues and conditions available in our panel. This analysis could be improved by including more transcriptomes in the future.

Using this approach, we find functional categories involving lncRNAs similar to those previously reported in both *A. thaliana* and other plant species. For example, we find 70 *A. thaliana* lncRNAs distributed in modules 5, 21, 18, and 39, all functionally enriched in coding genes associated with drought. Numerous lncRNAs are involved in this response in plants [129], including 664 lncRNAs in maize [130], 51 in cassava, 1096 and 126 in a drought-resistant variety of *Brassica napus* [131]. Similarly, we identified five modules with 72 lncRNAs related to response to pathogens (Fig. S12). lncRNAs have been previously found to be differentially expressed in response to infection in tomato [132] and maize [133].

Previous works have already established the relationship between lncRNAs and processes related to photosynthesis in *A. thaliana* and rice [134], as well as in the response to different types of light [18,135]. Photosynthesis is arguably the most important biological pathway in plants. Our results show that the function with the highest number of lncRNAs is related to chloroplast organization and response to light (Fig. S6); this indicates that a large number of lncRNAs may be involved in these processes. It is worth noting that most of our data were obtained from photosynthetic tissues and seedlings, which may explain why our largest modules, comprising the majority of lncRNAs, are associated with photosynthetic processes.

We also identified lncRNAs co-expressed with genes involved in root development and root response to multiple environmental stimuli (Fig. 4). lncRNAs have previously been shown to participate in root differentiation and response to different stress conditions in *A. thaliana* [16,120,136–138], *Medicago truncatula*, where 5561 lncRNAs change their expression in the root due to osmotic stress [139], and in *Populus*, where 295 lncRNAs change their expression during root development [117].

**Figure 4.**
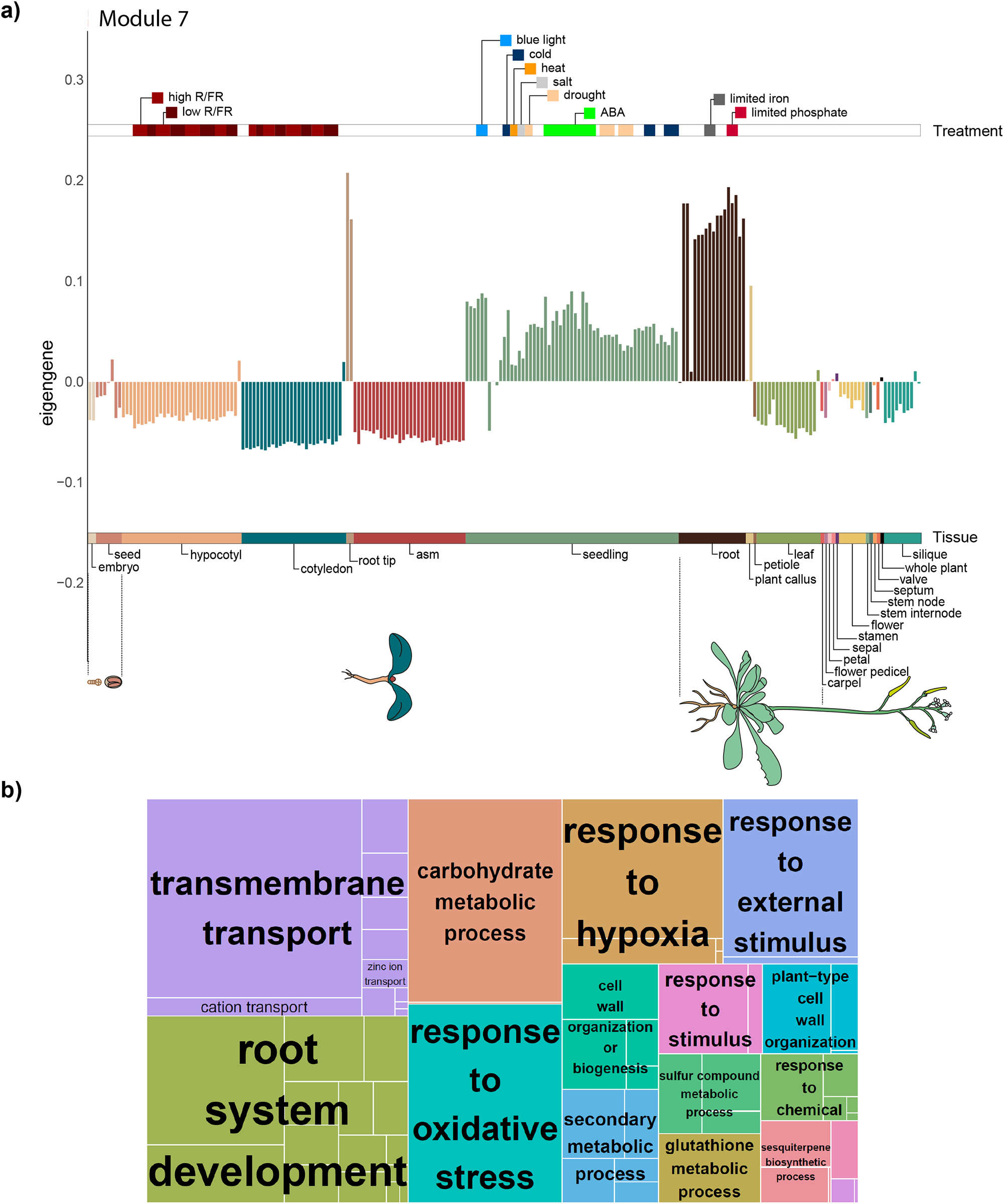
Module 7 expression and functions. a) Barplot of representative expression (eigengene) by transcriptome in module 7, which is enriched in genes expressed and participating in root development. Each bar represents the expression in one transcriptome, organized by tissue and ordered by age from younger (left) to older (right). The lower bar describes which tissue belongs to each color, while the upper one the different experimental treatments of each transcriptome. b) Treemap of enriched biological functions in module 7.

Surprisingly, several previously characterized lncRNAs, including *ELENA1, MARS, COOLAIR, IPS1*, and *HID1*, are not associated with any particular module. This might be partly because some of these lncRNAs perform their regulatory function in specific environmental conditions (e.g., prolonged cold in the case of *COOLAIR*), poorly represented in our transcriptomic panel [31]. Furthermore, the functional association via co-expression is not a fail-proof method; it only identifies lncRNAs expressed in most tissues sampled and that have a strong expression association with genes with similar functions. Thus, many other novel lncRNAs reported in our annotation with no functional association may have important functions that this approach could not identify.

We hope this highly curated, transcriptomic informed lncRNA annotation with functional associations via coexpression in *A. thaliana b*ecomes a valuable resource to the *A. thaliana* and the plant lncRNA community. In the future, we want to assess if the functional association relationships between lncRNAs and other RNAs are conserved in different species and how their loss or gain might be associated with the loss or gain of particular traits in this and other plant families.

## Declarations

### Ethics approval and consent to participate

Not applicable

### Consent for publication

Not applicable

## Availability of data and materials

Raw datasets, software, and documents are available under a CC-BY license at Github [140] and FigShare (see Supplementary Information) and NCBI (PRJNA765039).

## Competing interests

The authors declare that they have no competing interests

## Funding

This work was funded in part by Consejo Nacional de Ciencia y Tecnología (CONACYT Ph.D. Scholarships 338379 (JAC-G), 781634 (ELC-N), and 780678 (IJG-L) and by a Royal Society Newton Advanced Fellowship (NAF\R1\180303) awarded to SLF-V.

## Authors’ contributions

JAC-G and SLF-V conceived and coordinated the study. JAC-G made assembly and annotation of lncRNAs, as well as tissues specific analysis and coexpression analysis interpretation. ELC-N performed RNA-seq experiments and the identification and classification of lncRNAs. JAP-P helped with batch processing of RNA-seq data. IJG-L did the coexpression and functional enrichment analysis. SLF-V obtained the funding. SLF-V, JAC-G, ELC-N, and IJG-L drafted the manuscript. All authors read and approved the final manuscript.

## Acknowledgments

We acknowledge Dr. Katarzyna Oktaba for her library quality and preparation advice.

## Notes

### Competing Interest Statement

The authors have declared no competing interest.

