## Supplementary Information for "Transcriptome-guided annotation and functional classification of long non-coding RNAs in *Arabidopsis thaliana*"

### Supplementary Datasets

**Dataset S1** Name and position of all annotated lncRNAs in bed12 format with the biotype added in column number 13 (Private link: https://figshare.com/s/5d19c00590cbd7fe73a1)

**Dataset S2** Redundant lncRNAs biotypes in Bed12 format plus biotype in column number 13 (Private link: https://figshare.com/s/859d12a6da94ba62c9fa)

**Dataset S3** Table of metadata and ID of all transcriptomes used in this project (Private link: https://figshare.com/s/70c23951cda438d0fe35)

### Supplementary Tables

**Table S1 Information of root and shoot *A. thaliana* transcriptomes generated**

| Sample Name | SRA Accession Number | % GC | Total reads | Remaining reads after filtering |
|---|---|---|---|---|
| At_shootR1_1 | SRR16093081 | 46% | 36075409 | 31423073 |
| At_shootR1_2 | SRR16093081 | 47% | 36075409 | 31423073 |
| At_shootR2_1 | SRR16093080 | 46% | 49835624 | 43879678 |
| At_shootR2_2 | SRR16093080 | 47% | 49835624 | 43879678 |
| At_rootR1_1 | SRR16093079 | 45% | 40506616 | 35421322 |
| At_rootR1_2 | SRR16093079 | 45% | 40506616 | 35421322 |
| At_rootR2_1 | SRR16093078 | 44% | 40231170 | 35455572 |
| At_rootR2_2 | SRR16093078 | 45% | 40231170 | 35455572 |

**Table S2 lncRNAs with known functions present in a single tissue**

| Gene ID | lncRNA name | Tissue group |
|---|---|---|
| AT2G35747 | *HID1* | asm |
| AT1G08103 | *FLINC* | asm, carpel, flower |
| AT3G05945 | *AT3G05945* | embryo |
| AT4G08415 | *AT4G08415* | embryo |
| AT1G05987 | *AT1G05987* | flower |
| AT4G06195 | *AT4G06195* | hypocotyl, cotyledon |
| AT4G16355 | *ELENA1* | petal |
| AT2G34655 | *APOLO* | petiole |
| AT1G21529 | *DRIR* | plant callus |
| AT5G00580 | *MARS* | root |
| AT3G09922 | *IPS1* | root, plant callus |
| AT5G03545 | *AT4* | root, plant callus |
| AT5G01675 | *COOLAIR* | seed |
| AT5G06335 | *AT5G06335* | seedling, flower |
| AT5G06325 | *AT5G06325* | seedling, root |

### Supplementary Figures

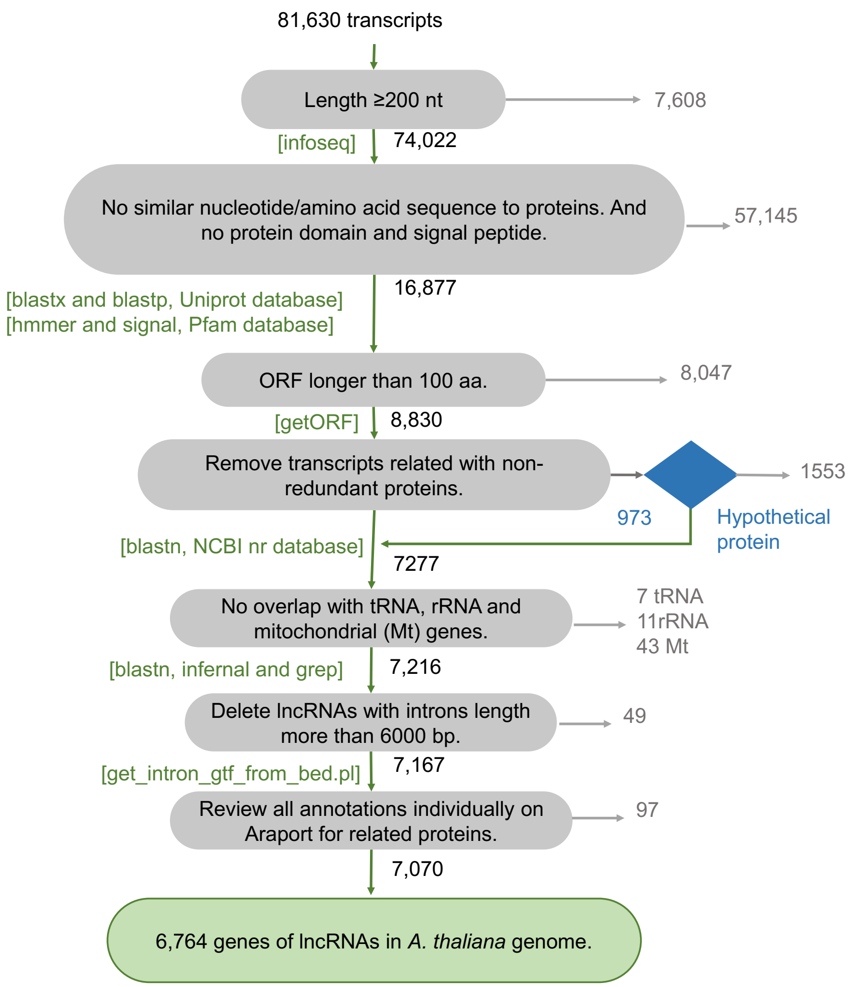


**Figure S1.** Diagram of filters used to annotate lncRNAs

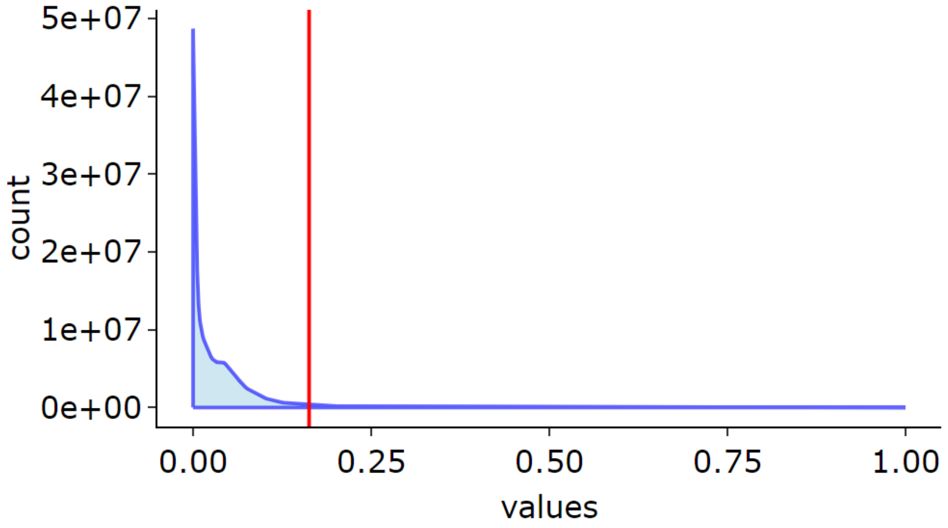


**Figure S2.** Cut-off point of SPM values by tissues

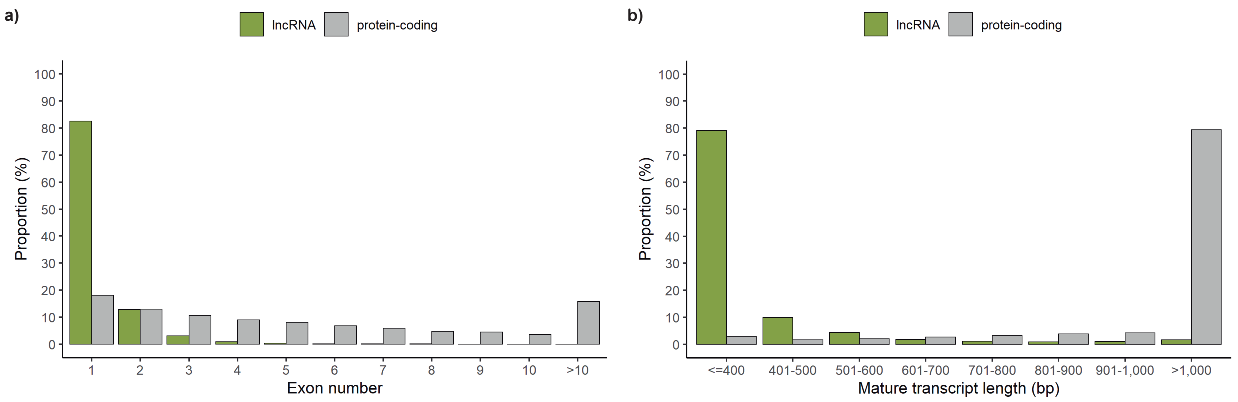


**Figure S3.** a) Proportion of the number of exons contained in the lncRNAs and coding genes. b) Proportion of the average size of the lncRNAs compared to the coding genes.

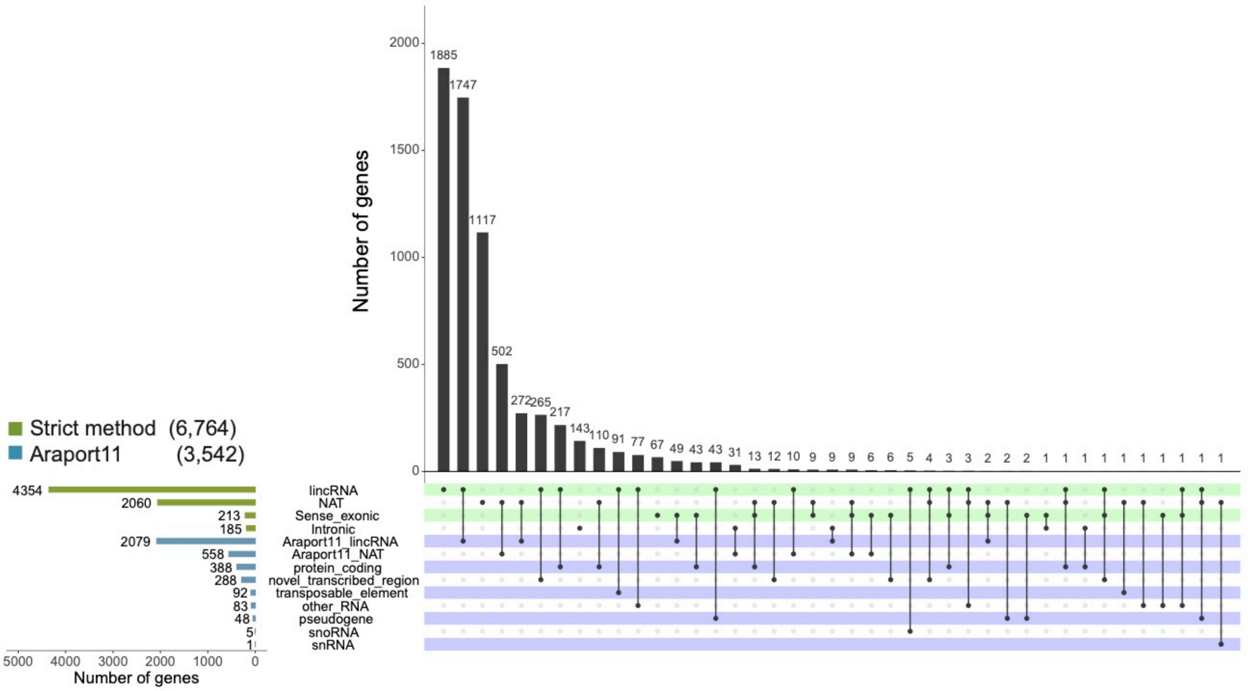


**Figure S4.** Comparison of each of the categories of lncRNAs present in Araport11 against the categories of the newly generated annotation.

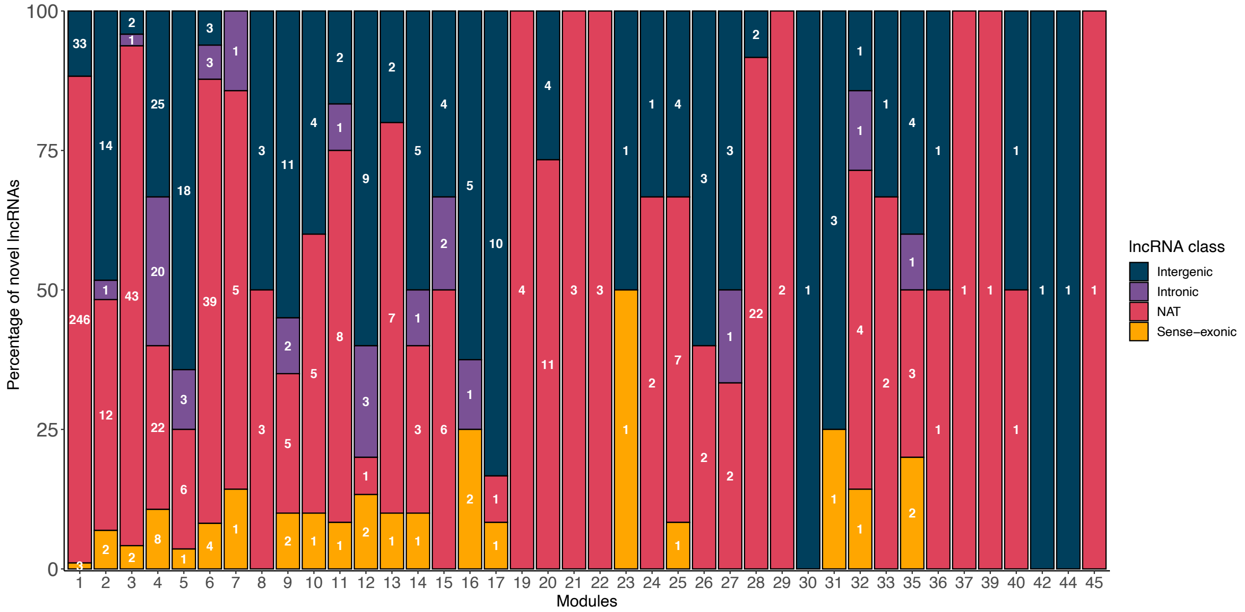


**Figure S5. Novel lncRNAs per module.** Proportion of novel lncRNAs (without annotation in Araport11) in each co-expression module and classified by biotype (Intergenic (lincRNA) - blue, Intronic - purple, NAT - pink, Sense-exonic - yellow).

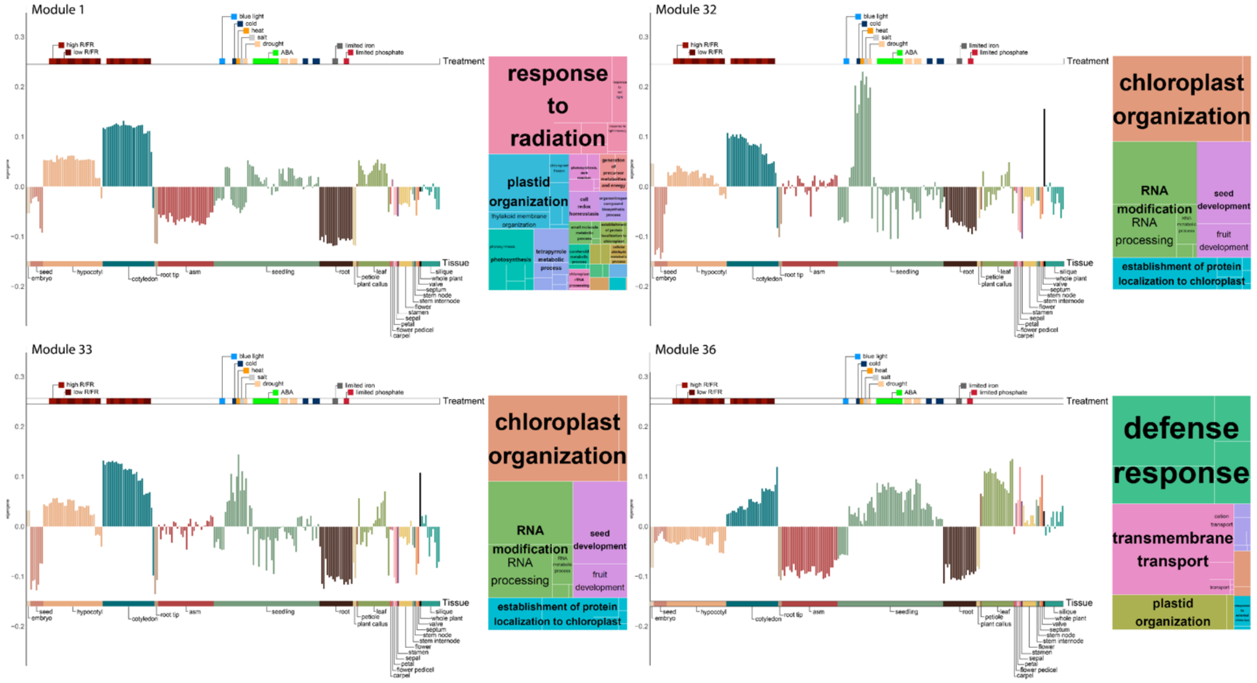


**Figure S6**. Eigengene expression per module for modules 1, 32, 33 and 36, chloroplast organization and photosynthesis functional category (4 modules with 409 lncRNAs)

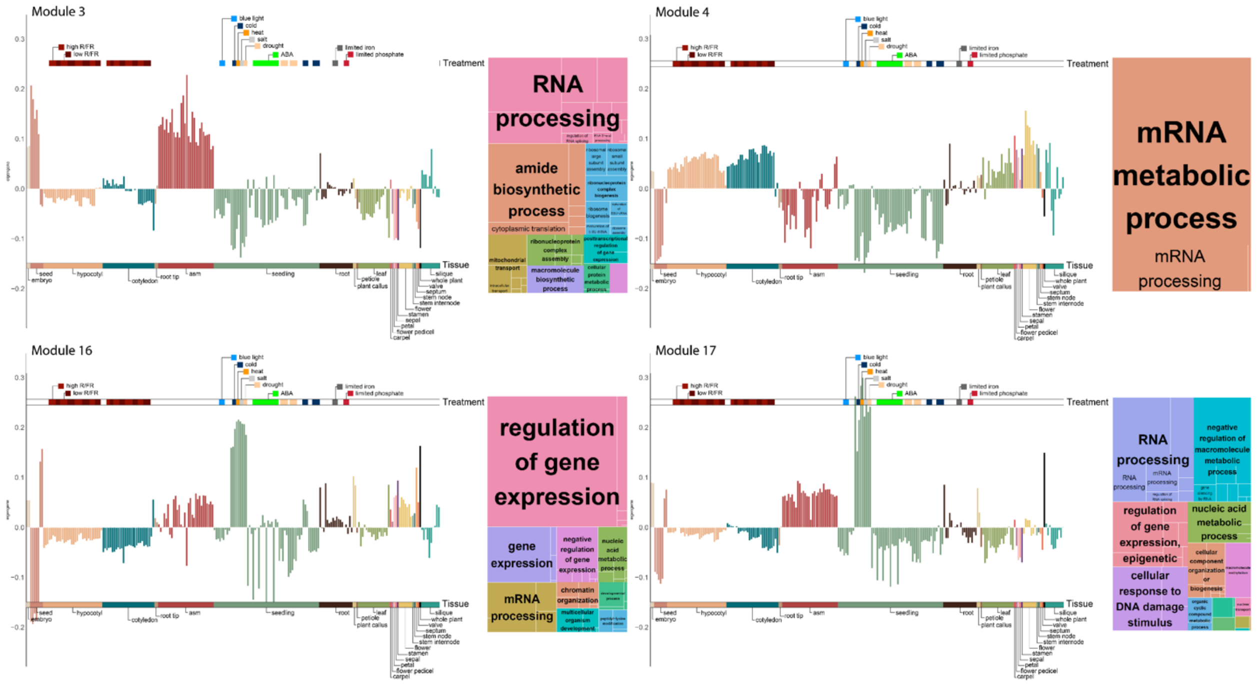


**Figure S7**. Eigengenes expression per module for modules 3, 4, 16 and 17; RNA regulation and transcription functional category (4 modules with 375 lncRNAs)

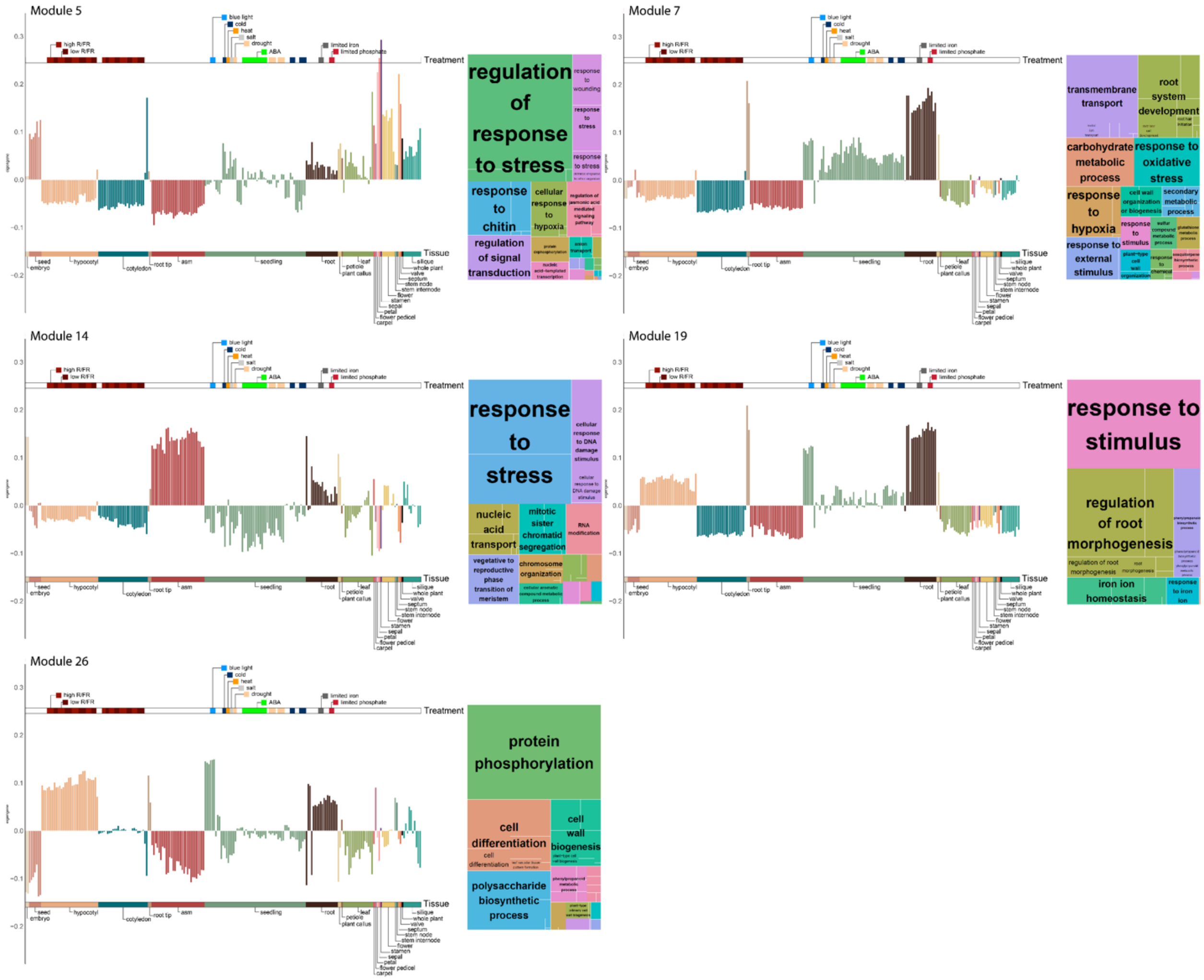


**Figure S8**. Eigengene expression per module for modules 5, 7, 14, 19 and 26; root development and response to root-related stress  functional category (5 modules with 125 lncRNAs)

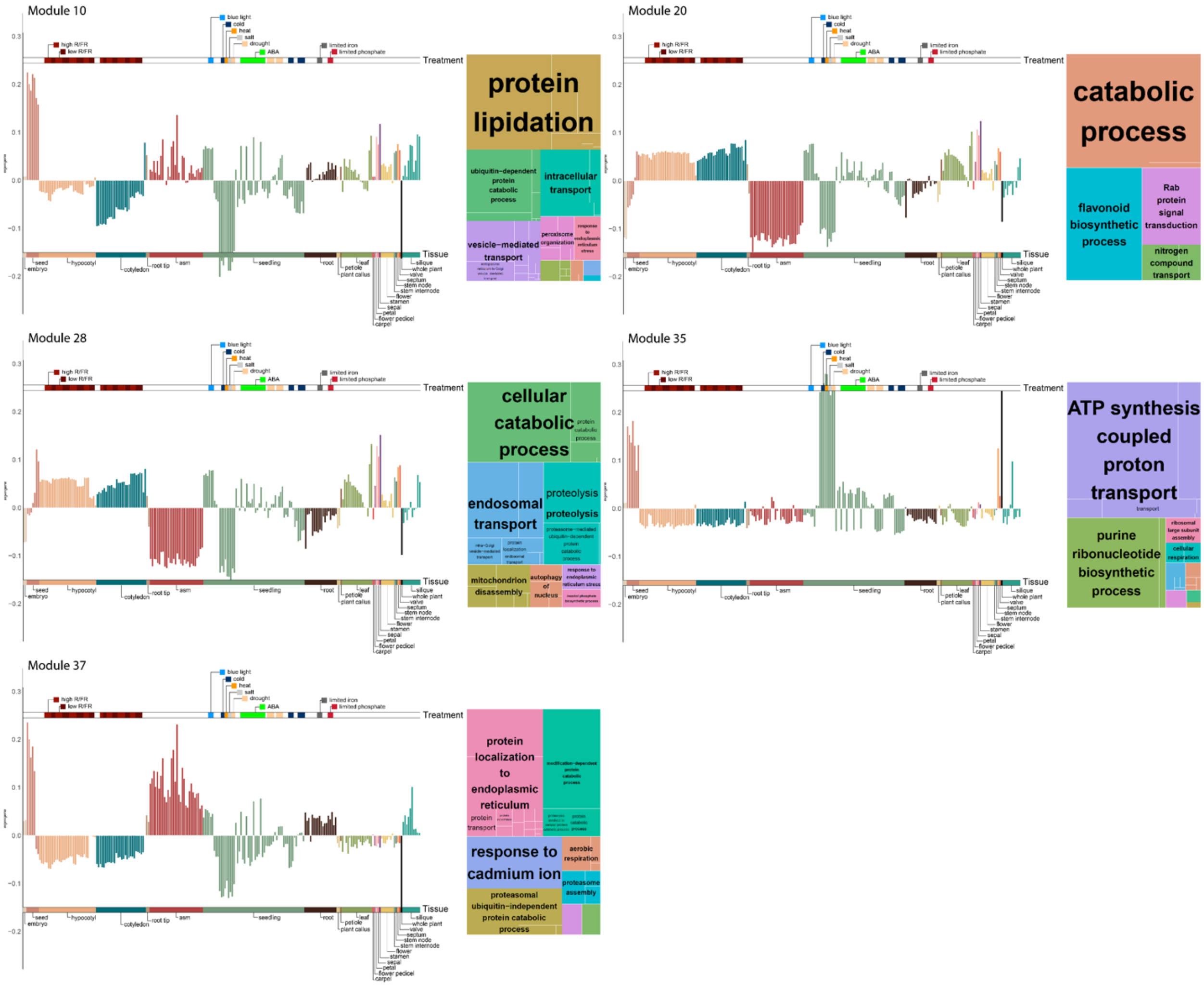


**Figure S9**. Eigengene expression per module for modules; protein labeling and transport functional category (5 modules with 117 lncRNAs)

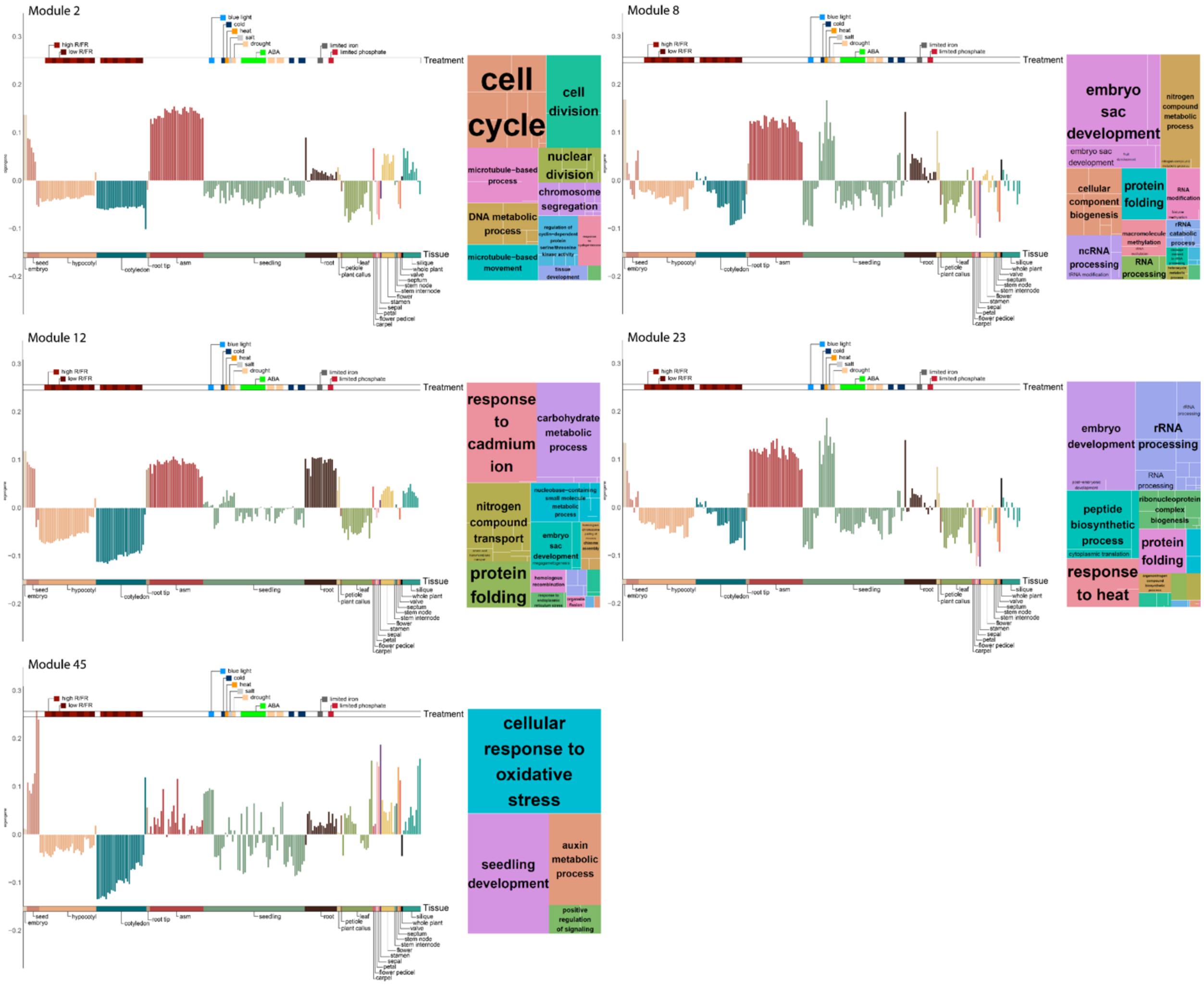

**Figure S10**. Eigengene expression per module for modules 2, 8, 12, 23 and 45; cell division functional category (5 modules with 112 lncRNAs)

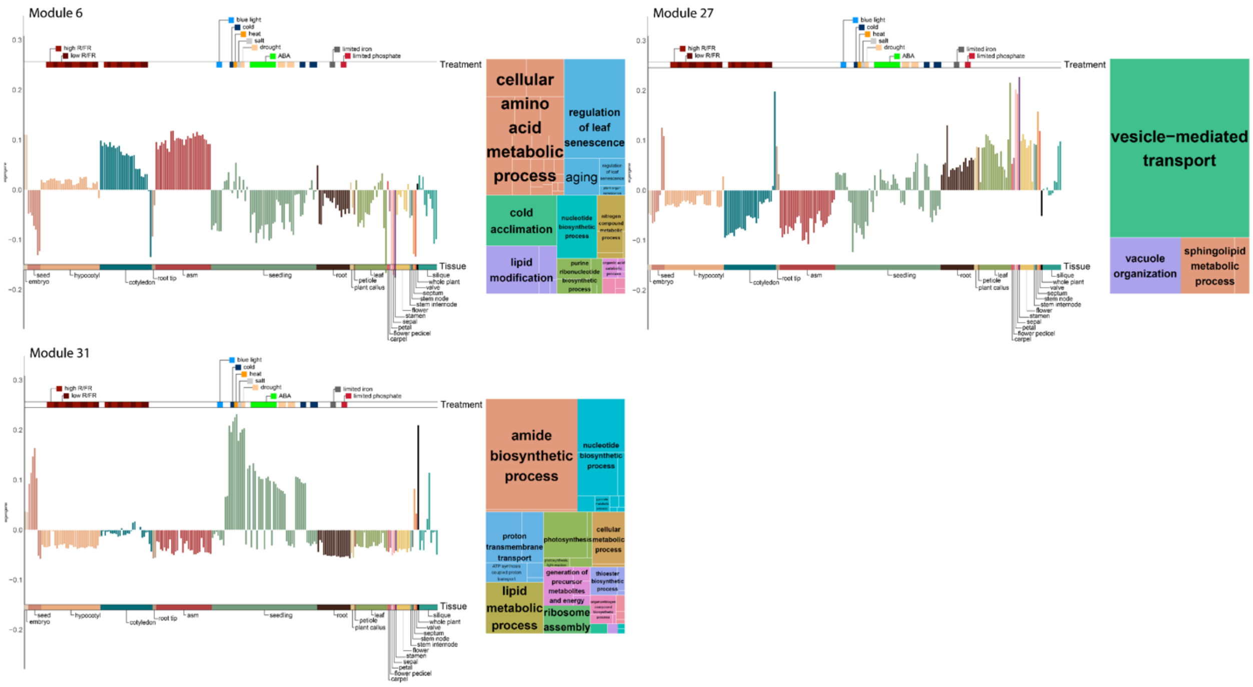


**Figure S11**. Eigengene expression per module for modules 6, 27 and 31; lipids and membranes functional category (3 modules with 97 lncRNAs)

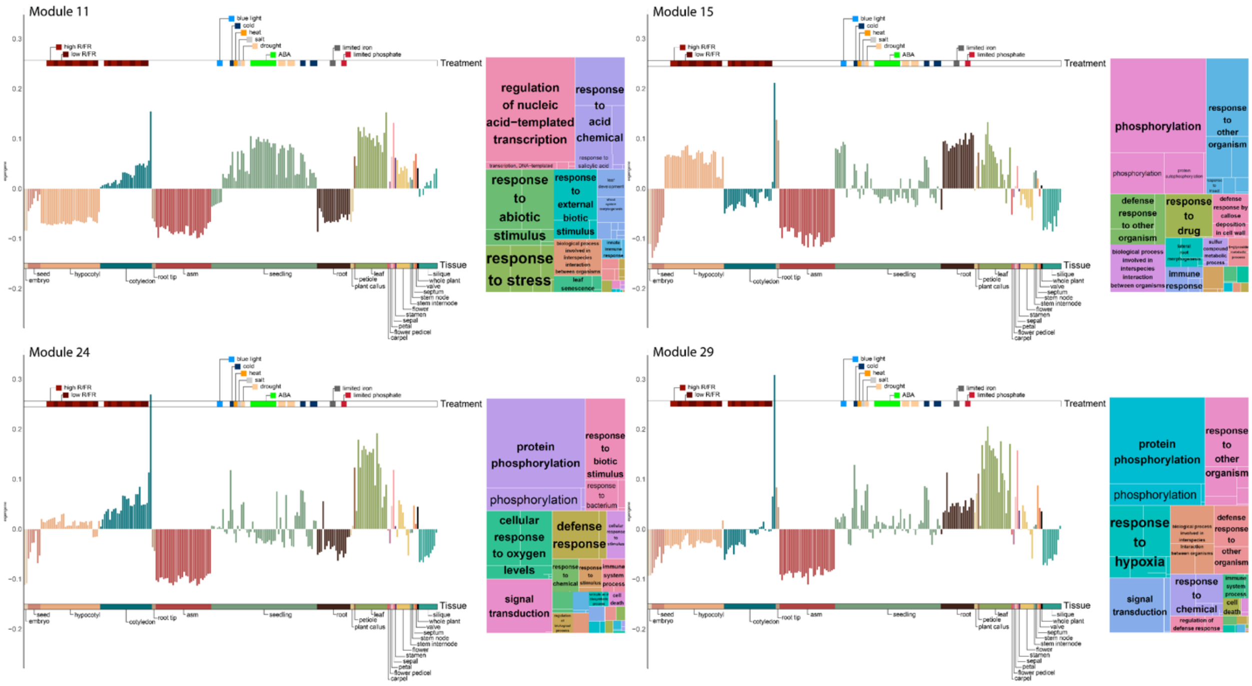


**Figure S12**. Eigengene expression per module for modules 11, 15, 24 and 29; response to pathogens functional category (4 modules with 72 lncRNAs)

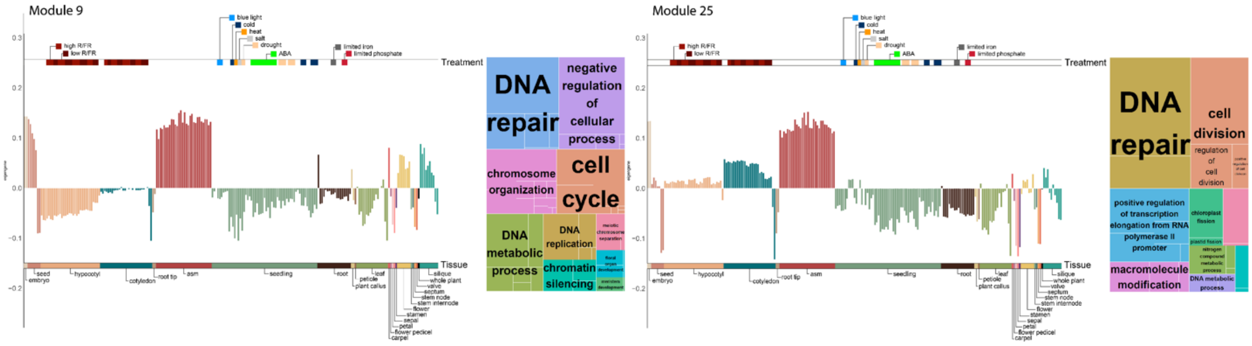


**Figure S13**. Eigengene expression per module for modules 9 and 25; DNA repair functional category (2 modules with 61)

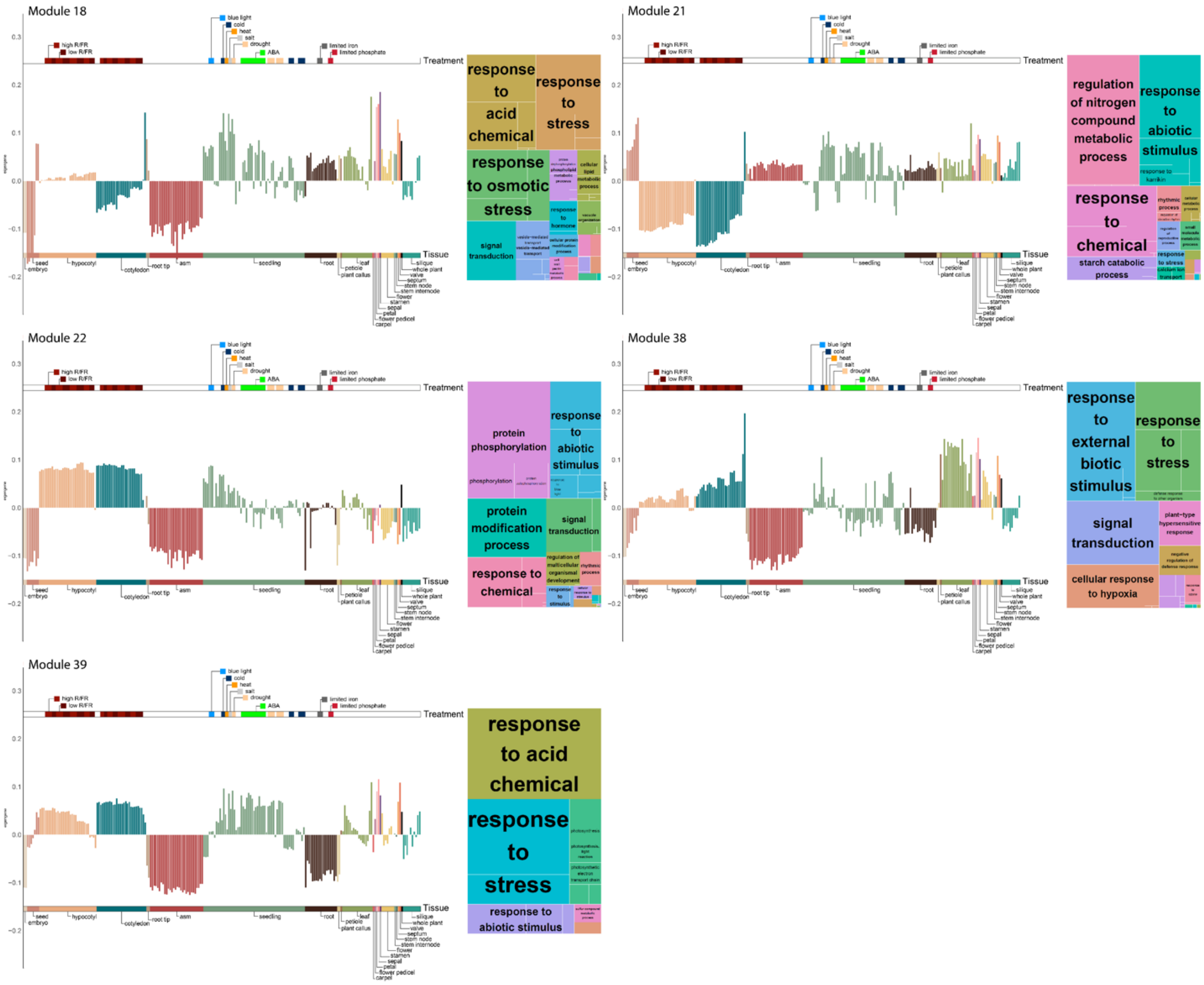


**Figure S14**. Eigengene expression per module for modules 18, 21, 22, 38 and 39; response to stress functional category (5 modules with 17 lncRNAs)
